# Forest tree extracts induce resistance to *Pseudomonas syringae* pv. *tomato* in Arabidopsis

**DOI:** 10.1101/2023.10.24.563420

**Authors:** Veedaa Soltaniband, Adam Barrada, Maxime Delisle-Houde, Martine Dorais, Russell J. Tweddell, Dominique Michaud

**Author notes:** E-mail addresses: < >, < >, < > < >, < > < >. Correspondence: Dominique Michaud Russell J. Tweddell.

## Abstract

**Background:** The widespread use of conventional pesticides to control plant fungal and bacterial pathogens poses significant risks to human health and the environment, and there is an urgent need for safer and more sustainable alternatives in agricultural management. Studies have shown that plant extracts can be effective in controlling plant diseases either by directly targeting the pathogens or by reinforcing the host plant’s own defenses. Here, we examined the potential of ethanolic extracts from forest tree species eastern hemlock, English oak, eastern red cedar and red pine for their antibacterial activity against *Pseudomonas syringae* pv. *tomato* (*Pst*) strain DC3000 and the ability of these forestry by-products to trigger effective defense responses in the model plant *Arabidopsis thaliana*.

**Results:** The four tree extracts exhibited direct toxic effects against *Pst* DC3000, as notably observed for the English oak extract inhibiting bacterial growth and showing bactericidal activity at relatively low concentrations. Using an Arabidopsis line expressing reporter protein ß-glucuronidase under the control of a salicylic acid-inducible pathogenesis-related protein gene promoter, the extracts were shown also to induce defense-related genes expression in leaf tissue. RT-qPCR assays with DNA primers for different gene markers further confirmed this conclusion and highlighted gene-inducing effects for the tree extracts triggering, at different rates, the expression of salicylic acid- and oxidative stress-responsive genes. The extracts direct antibacterial effects, combined with their defense gene-inducing effects *in planta*, resulted in a strong host plant-protecting effect against *Pst* DC3000 associated with bacterial growth rates reduced by ∼75 to 98% seven days post-infection, depending on the extract.

**Conclusions:** These findings show the effectiveness of tree extracts as eventual plant protectants against the plant bacterial pathogen *Pst*. In a broader perspective, they suggest the potential of these forestry by-products as a source of bioactive compounds useful in plant protection and as a sustainable, eco-friendly alternative to conventional synthetic pesticides for the management of economically important plant pathogens.

## Background

Plant diseases pose a significant threat to agricultural and horticultural crops worldwide, resulting in substantial losses in crop productivity, quality and economic value [1]. The use of conventional pesticides, such as fungicides and bactericides, is widespread for controlling plant pathogenic fungi and bacteria [2,3]. However, their extensive use has faced increasing criticism due to the various problems they cause to agriculture, human health and the environment [4,5,6]. It is therefore crucial to urgently develop environmentally friendly, human-safe alternatives to conventional pesticides that offer a sustainable approach to plant pathogen control.

Over the past two decades, several studies have demonstrated the potential of plant extracts in limiting the development of plant diseases and highlighted their potential as a sustainable alternative to conventional pesticides [7,8,9,10,11]. Numerous studies have reported the efficacy of such extracts to prevent bacterial and fungal diseases on cultivated plants, caused notably by fungal pathogens such as *Puccinia triticina*, *Magnaporthe grisea*, *Fusarium oxysporum* f. sp. *lycopersici* and *Botrytis cinerea*, or by bacterial pathogens such as *Pseudomonas cichorii*, *Xanthomonas campestris* pv. *vitians* and *Xanthomonas vesicatoria* [12,13,14,15,16,17]. The control of microbial diseases using plant extracts has in several cases been explained by their direct toxic effects on the pathogens [12,17,18]. It has also been associated with the induction of plant natural defenses leading to the accumulation of antimicrobial proteins and organic toxic compounds in host plant tissues [13,14,15,19].

Here, we investigated the plant protective potential of ethanolic extracts from twigs or leaves of forest tree species eastern hemlock (*Tsuga canadensis*), eastern red cedar (*Juniperus virginiana*), English oak (*Quercus robur*) and red pine (*Pinus resinosa*). Studies were conducted both to monitor the direct antibacterial activity of these forestry by-products, and to assess their ability to activate host plant natural defenses eventually detrimental to plant pathogens. We used as a pathosystem the model plant *Arabidopsis thaliana* infected with *Pseudomonas syringae* pv. *tomato* strain DC3000 (*Pst* DC3000) [20,21], the causal agent of tomato bacterial speck in tomato cultivation setups worldwide [22].

## Results and discussion

### Forest tree extracts are toxic to *Pst* DC3000

Minimum inhibitory concentrations (MIC) and minimum bactericidal concentrations (MBC) were first determined to evaluate the direct toxic effects of the extracts against *Pst* DC3000 (**Table 1**). MIC values, that refer to the lowest concentrations of extracts at which no metabolic activity of *Pst* DC3000 was observed following a 24-h incubation period, ranged from 0.8 to >25 mg mL^-1^ from one extract to another, with English oak extract showing the lowest value (at 0.8 mg mL^-1^) and hence the strongest antibacterial effect. MBC values, that refer to the lowest concentrations of extracts killing ≥99.9% of the bacteria following a 24-h incubation period, ranged from 25 to more than 50 mg mL^-1^. Again, the English oak extract was the most potent, with an MBC of 25 mg mL^-1^ compared to MBC values greater than 50 mg mL^-1^ for the other three extracts. As expected, given the negligible direct toxic effects of this compound [23], salicylic acid (SA) functional analogue benzothiadiazole (BTH) (commercialized as ACTIGARD^TM^ 50WG) used as a positive control for SA-inducible defense gene inductions (see below) showed no obvious toxic effect against the bacterium, with MIC and MBC values both greater than 50 mg mL^-1^.

**Table 1.** Minimum inhibitory concentrations (MIC) and minimum bactericidal concentrations (MBC) of tree extracts against bacterial pathogen *P. syringae* pv. *tomato* strain DC3000.

| Tree extract | MIC (mg mL<sup>-1</sup>)<sup>1</sup> | MBC (mg mL<sup>-1</sup>)<sup>1</sup> |
| --- | --- | --- |
| Eastern hemlock (*Tsuga canadensis*) | 25 | >50 |
| Eastern red cedar (*Juniperus virginiana*) | 12.5 | >50 |
| English oak (*Quercus robur*) | 0.8 | 25 |
| Red pine (*Pinus resinosa*) | 12.5 | >50 |
| BTH (ACTIGARD™ 50WG)<sup>2</sup> | >50 | > 50 |

<sup>1</sup> Each value is the mean of six replicates.
<sup>2</sup> SA functional analogue used as a negative, non-toxic control for the toxicity assays.

Previous studies have reported the pharmacological and therapeutic benefits of forest tree extracts in animals and humans, notably including the extracts of English oak and related species of the *Quercus* genus [24,25,26]. In complement, our findings suggest the potential of such extracts as antibacterial phytosanitary agents in crop protection, similar to Cibele et al. [27] reporting the antibacterial activity of aqueous extracts from twelve Brazilian medicinal plants against the plant pathogens *Acidovorax citrulli*, *Pectobacterium carotovorum* subsp. *carotovorum*, *Ralstonia solanacearum* and *X. campestris* pv. *campestris*. Several secondary organic compounds in plants exhibit antimicrobial activity, such as for instance various terpenes, alkaloids and flavonoids found in red pine, English oak and other forest trees [28]. Studies will be welcome in coming years to identify the antibacterial determinants of the tree extracts considered in this study, notably for the English oak extract showing relatively low MIC and MBC values.

### Tree extracts induce SA-inducible genes in Arabidopsis

Defense gene-inducing effects of the tree extracts were assessed using pathogenesis reporter line PR1::GUS [29], a transgenic Arabidopsis line engineered to express reporter protein ß-glucuronidase (GUS) under the control of *PR1* promoter, an SA-inducible promoter driving the expression of pathogenesis-related protein 1 (PR1) upon biotroph or hemibiotroph pathogen infection (**Fig. 1**). GUS expression in the PR1::GUS line was induced by all extract treatments, albeit at different levels depending on the extract. GUS staining was observed in all organs considered, including the cotyledons, the hypocotyl and the true leaves (**Fig. 1A**). Eastern hemlock, eastern red cedar and red pine extracts exhibited strong *PR1* promoter-inducing effects compared to the negative control treatment, roughly comparable to the inducing effect of SA functional analogue BTH used as a positive control for SA-inducible *PR1* gene expression [30]. By comparison, plants treated with the English oak extract showed a faint GUS staining in leaf tissues, indicating a weaker promoter-inducing effect compared with the other three extracts (**Fig. 1A,B**).

**Fig. 1.**
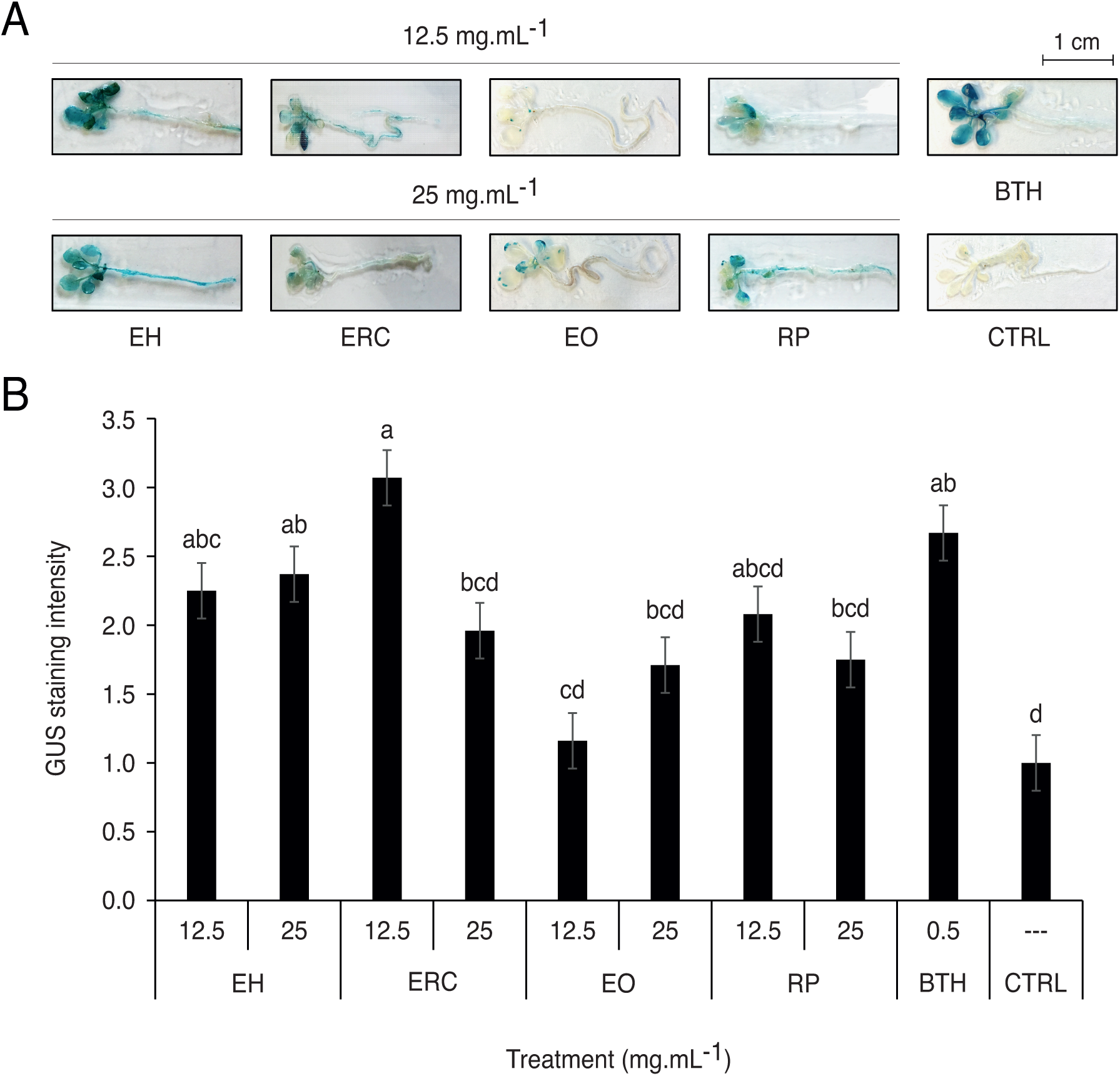
*PR1* gene induction in transgenic Arabidopsis PR1::GUS line treated with eastern hemlock (EH), eastern red cedar (ERC), English oak (EO) or red pine (RP) extracts. **A**, Representative seedling images showing GUS staining following plant extract treatment; **B**, GUS total intensity per plant after treatment with plant extracts at working concentrations of 12.5 and 25 mg mL^−1^, or with BTH (ACTIGARD^TM^ 50WG) at 0.5 mg mL^−1^. Data are presented relative to control seedlings treated with sterile water (relative value of 1.0). Each value is the mean of three replicates ± SE. Means with the same letter are not significantly different (post-ANOVA LSD, with an alpha threshold value of 5%).

RT-qPCR assays were carried out for *PR1* and additional SA-inducible genes (**Table 2**) to confirm the differential inducing effects of tree extracts on endogenous *PR1* gene promoter activity (*see* **Fig. 1**) and to highlight a possible induction of the SA signaling pathway leading to systemic acquired resistance (SAR) and eventual host plant resistance to pathogenic attack [31,32,33,34] (**Table 3**). DNA primers were also used for *NPR1*, the DNA sequence of master regulator NONEXPRESSOR OF PATHOGENESIS-RELATED GENES 1 (NPR1) protein [35,36]. As expected, given the limited effect of SA on NPR1 expression in Arabidopsis [37,38], BTH had little effect and the four extracts a variable, relatively limited impact on the abundance of *NPR1* transcripts. By comparison, the extracts strongly induced the transcription of endogenous *PR1* in treated plants relative to the control, similar to BTH and in accordance with their inducing effects on GUS expression in the *PR1::GUS* line (**Fig. 1**). The *WRKY70* gene, coding for SA signaling-inducing transcription factor WRKY70 [39], was also induced at various rates by BTH and the extracts, similar to SA-inducible genes for antimicrobial proteins GLUCAN ENDO-1,3-BETA-D-GLUCOSIDASE (PR2) [40] and KUNITZ TRYPSIN INHIBITOR 4 (KTI4, also referred to as KTI1) [41] produced downstream along the SA defense pathway. For instance, transcript numbers were increased by up to 7.8-fold for *WRKY70* in BTH-treated plants, or by 7.4-fold for *KTI4* and 3.4-fold for *PR2* in plants treated with the eastern red cedar and eastern hemlock extracts, respectively.

**Table 2.**
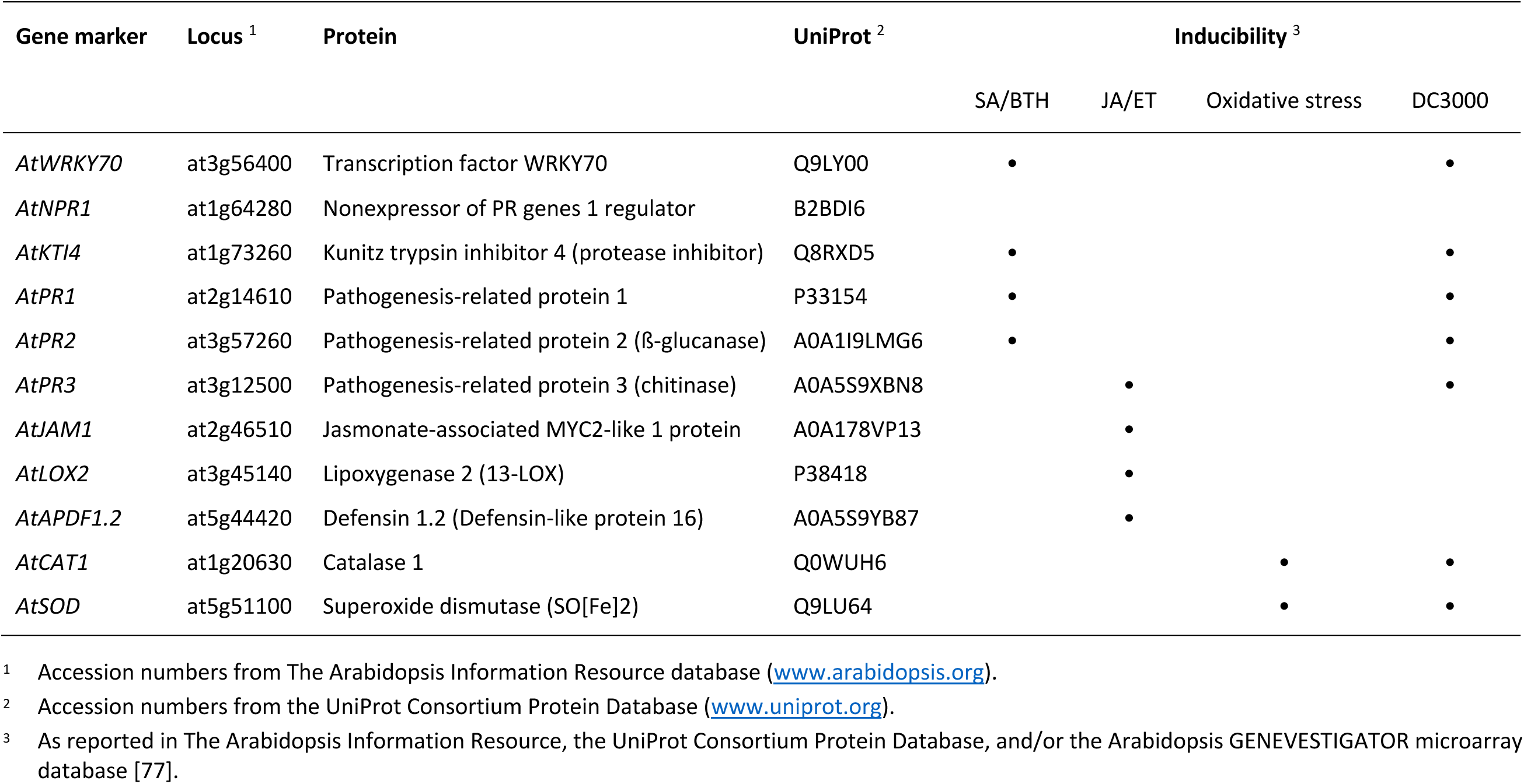
Marker genes selected for the qPCR analyses.

**Table 3.**
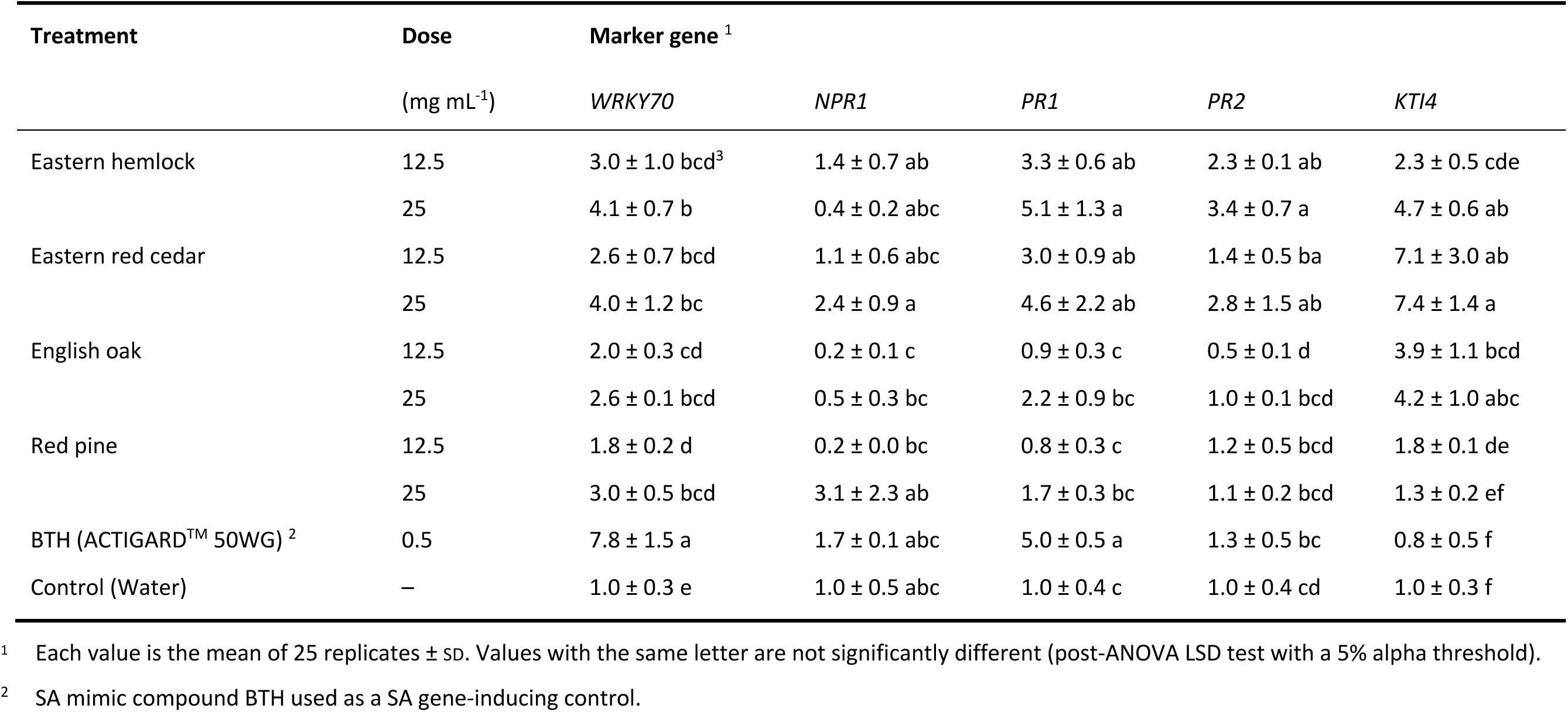
Relative expression of SA-inducible marker genes in plants responding to forest tree by-product extracts.

In agreement with the differential effects of plant extracts on GUS expression, the gene inducing effects were variable from one extract to another, from a null, even negative effect for some extracts used at low concentration to more than 100% increases in 25 extract dose–gene combinations out of 40 combinations assessed (**Table 3**). Also in line with the GUS expression assay, endogenous *PR1* transcriptional induction by the English oak and the red pine extracts, with transcript levels 0.8-to 2.2-fold compared to the control treatment, were weaker than the 3.0-to 5.1-fold inductions observed for the eastern hemlock and eastern red cedar extracts. Variable inductions were observed not only between the plant extracts for a given marker gene, but also among the marker genes for a given plant extract. For instance, the eastern red cedar extract induced *NPR1* by 10% and *PR2* by 40% at the lowest dose applied, compared to an induction rate of 710% for *KTI4* at the same dose. Likewise, *PR1* expression was increased by 70% in plants treated with the highest dose of red pine extract, lower than a 200% increase observed for *WRKY70* with the same treatment.

Several studies have reported the induction of PR protein-encoding and other SA-inducible genes in leaf tissue following treatment with plant extracts, as exemplified with crude extracts of seaweed [42,43,44], red grape [45], African lily [19], coffee [14] or medicinal plants [46]. For instance, Goupil et al. [45] reported the local and systemic induction of SA-inducible proteins PR1 and PR2 in tobacco leaves treated with an aqueous extract of red grape, similar to Islam et al. [43] reporting the induction of *PR1* and *NPR1* marker genes in Arabidopsis following foliar application of a seaweed extract; or to Medeiros et al. [14] showing an increased expression of WRKY family transcription factors involved in SAR in tomato treated with a coffee leaf extract. Overall, our observations confirmed that ethanolic extracts from forest tree by-products, such as those considered in this study, could trigger the transcription of SA-inducible genes including *WRKY70* for the biosynthesis of WRKY70, a key inducer of the SAR response leading to resistance against *Pst* DC3000 in Arabidopsis [47]. On a broader basis, the upregulation of SA-inducible antimicrobial proteins, such as PR2 and KTI4 in this study, suggested the potential of these extracts to promote the accumulation of defense compounds in host plant tissues eventually detrimental to biotrophic or hemibiotrophic pathogenic invaders.

### The tree extracts also induce SA-independent stress-related genes in leaf tissue

RT-qPCR assays were conducted for marker genes induced by the SA antagonist jasmonic acid (JA) [48] and marker genes induced under oxidative stress conditions (see **Table 2**) to confirm the specificity of the tree extracts in triggering SA-inducible genes induction or, on the opposite, to highlight broader gene-inducing effects eventually promoting concurrent stress-related metabolic routes (**Table 4**). As expected, BTH showed no inducing effect on the transcription of JA-inducible genes *JAM1*, *LOX2, PR3* and *PDF1.2* compared to control plants, even showing a marginal downregulating effect on the four genes likely explained by the well characterized antagonistic effects of SA [49,50], NPR1 [51] and WRKY70 [47,52] on the JA defense pathway. By contrast, the four tree extracts showed a significant inducing effect on the transcription of *PR3*, a gene marker for JA [and ethylene] signaling induced by different fungal and bacterial pathogens [53]. More specifically, extract-treated plants showed *PR3* transcript levels 2 to 7 times greater than in water-treated plants, and 3 to 10 times greater than in BTH-treated plants, at the highest dose of extract. Considering the previously reported inducing effect of JA and null effect of SA on *PR3* expression [54,55], these observations suggested the occurrence of additional defense gene-inducing triggers in the extracts acting along with, but independently of, the SA signaling pathway in leaf tissue. Activation of *PR3* by the tree extracts was likely independent of the JA signaling pathway given the strong induction of SAR-inducible genes in leaves and the mutual antagonistic relationship generally established *in planta* between SA and JA. Dual induction of the SA and JA signaling pathways promoting host resistance to *Pst* DC3000 has been observed previously in Arabidopsis leaves presenting an effector-triggered immunity hypersensitive response [56] or treated with the polysaccharide elicitor chitosan [57]. By comparison, the weak, marginal negative effects here observed for BTH on JA-inducible genes *JAM1*, *LOX2*, *PDF1.2* and *PR3* suggested instead an inducing scheme independent of the SA and JA signaling pathways for PR3 protein production.

**Table 4.** Relative expression of JA/ET and oxidative stress-marker genes in plants responding to forest tree by-product extracts.

| Treatment | Dose<br>(mg mL<sup>-1</sup>) | Marker gene <sup>1</sup> |  |  |  |  |  |
| --- | --- | --- | --- | --- | --- | --- | --- |
|  |  | *JAM1* | *LOX2* | *PR3* | *PDF1.2* | *CAT1* | *SOD* |
| Eastern hemlock | 12.5 | 0.7 ± 0.1 bcd<sup>4</sup> | 1.2 ± 0.2 b | 1.8 ± 0.3 bc | 0.9 ± 0.1 abc | 1.9 ± 0.3 cd | 0.9 ± 0.3 cd |
|  | 25 | 0.9 ± 0.2 bcd | 1.8 ± 0.3 ab | 3.0 ± 0.5 b | 0.9 ± 0.4 abc | 3.0 ± 0.8 bc | 1.3 ± 0.2 abc |
| Eastern red cedar | 12.5 | 1.2 ± 0.3 a | 0.9 ± 0.1 b | 7.3 ± 1.7 a | 1.6 ± 0.7 abc | 282.0 ± 55.0 a | 0.6 ± 1.1 ab |
|  | 25 | 1.2 ± 0.3 ab | 1.3 ± 0.2 b | 6.8 ± 1.8 a | 1.7 ± 0.6 ab | 423.0 ± 86.0 ab | 0.6 ± 0.3 a |
| English oak | 12.5 | 0.6 ± 0.2 cd | 0.9 ± 0.2 b | 2.3 ± 0.5 b | 0.5 ± 0.1 c | 2.0 ± 0.6 cd | 2.5 ± 0.0 cd |
|  | 25 | 0.4 ± 0.2 d | 0.9 ± 0.3 b | 2.1 ± 0.5 b | 2.3 ± 0.9 a | 22.0 ± 20.0 ab | 2.3 ± 0.2 d |
| Red pine | 12.5 | 0.6 ± 0.1 cd | 4.3 ± 1.0 a | 1.6 ± 0.3 bc | 1.9 ± 0.6 ab | 0.9 ± 0.1 e | 0.8 ± 0.1 cd |
|  | 25 | 1.2 ± 0.2 ab | 1.0 ± 0.2 b | 2.7 ± 0.4 b | 0.8 ± 0.1 abc | 1.5 ± 0.3 cde | 0.7 ± 0.2 cd |
| BTH (ACTIGARD™ 50WG) <sup>2</sup> | 0.5 | 0.7 ± 0.2 abcd | 0.8 ± 0. b | 0.7 ± 0.1 d | 0.9 ± 0.2 bc | 1.3 ± 0.2 de | 1.0 ± 0.2 bcd |
| Control (water) | — | 1.0 ± 0.1 abc | 1.0 ± 0.3 b | 1.0 ± 0.3 cd | 1.0 ± 0.3 abc | 1.0 ± 0.0 e | 1.0 ± 0.3 cd |

<sup>1</sup> Each value is the mean of 25 replicates ± sd. Values with the same letter are not significantly different (post-ANOVA LSD test with a 5% alpha threshold).
<sup>2</sup> SA mimic compound BTH used as a SA gene-inducing control.

An alternative route could also explain the strong positive impact of some tree extracts on the expression of oxidative stress marker gene *CAT1* (**Table 4**). Compared with water- and BTH-treated plants, *CAT1* transcripts were increased by 3-fold to several orders of magnitude in leaves treated with the extracts of eastern hemlock, English oak and eastern red cedar. Noticeably, this inducing effect led to *CAT1* transcript levels increased by more than 400 times for the cedar extract compared to control and BTH-treated plants. The basic cause of *CAT1* induction remains unknown at this point but a plausible trigger could be hydrogen peroxide (H_2_O_2_), the substrate of CAT1, possibly generated in leaves upon extract treatment. Forest trees produce an array of secondary compounds to defend themselves against arthropods and pathogens [28], that can not only be detrimental to microbial pathogens as shown above for *Pst* DC3000 (see **Table 1**) but also induce the production of secondary messengers like H_2_O_2_ and other reactive oxygen species (ROS) involved in acclimation to abiotic stress conditions [58,59]. For instance, terpenoids present in large amounts in conifer (e.g., cedar) and other plant extracts show promise as potential biopesticides in agriculture for pest and pathogen control [60]. Considering the cell membrane-destabilizing and ROS (e.g., H_2_O_2_)-generating effects reported for terpenoids in plant extracts [61], these compounds could have here triggered the induction of oxidative stress-inducible genes, such as *CAT1*, in tree residue-treated leaves. In support to this hypothesis, H_2_O_2_ is required for *CAT1* expression in Arabidopsis leaves [62], the CAT1 enzyme is readily induced by exogenous H_2_O_2_ treatment and abiotic stress cues (e.g., drought, abscisic acid, salts) that promote H_2_O_2_ production in Arabidopsis [62], and the general role of stress-inducible catalases in plants is the elimination of excess H_2_O_2_ accumulated in leaves under stress conditions [63].

### Tree extract-treated plants are resistant to *Pst* DC3000

Together, our observations pointed to broad defense/stress-related gene inducing effects for the tree extracts in Arabidopsis leaves involving, on the one hand, an effective activation of the SAR response and, on the second hand, the induction of additional defense- or oxidative stress-inducible genes, dependent on the specific chemical composition of each extract. Numerous studies have discussed the practical potential of SA mimics like BTH and other functional analogues to induce a SAR response leading to biotroph and hemibiotroph pathogen resistance in plants of economic importance [64]. Similarly, several studies have reported resistance to pathogenic fungi or bacteria in plants treated with SAR-inducing plant extracts or products (e.g., [42,46,65]) or in plants genetically engineered to express SAR-inducing regulators such as NPR1 or WRKY70 [47,66,67]. As a complement, bioassays were here carried out to assess the protective potential of the four tree extracts against bacterial pathogens, again using as a model the Arabidopsis–*Pst* DC3000 pathosystem (**Figs. 2** and **3**).

**Fig. 2.**
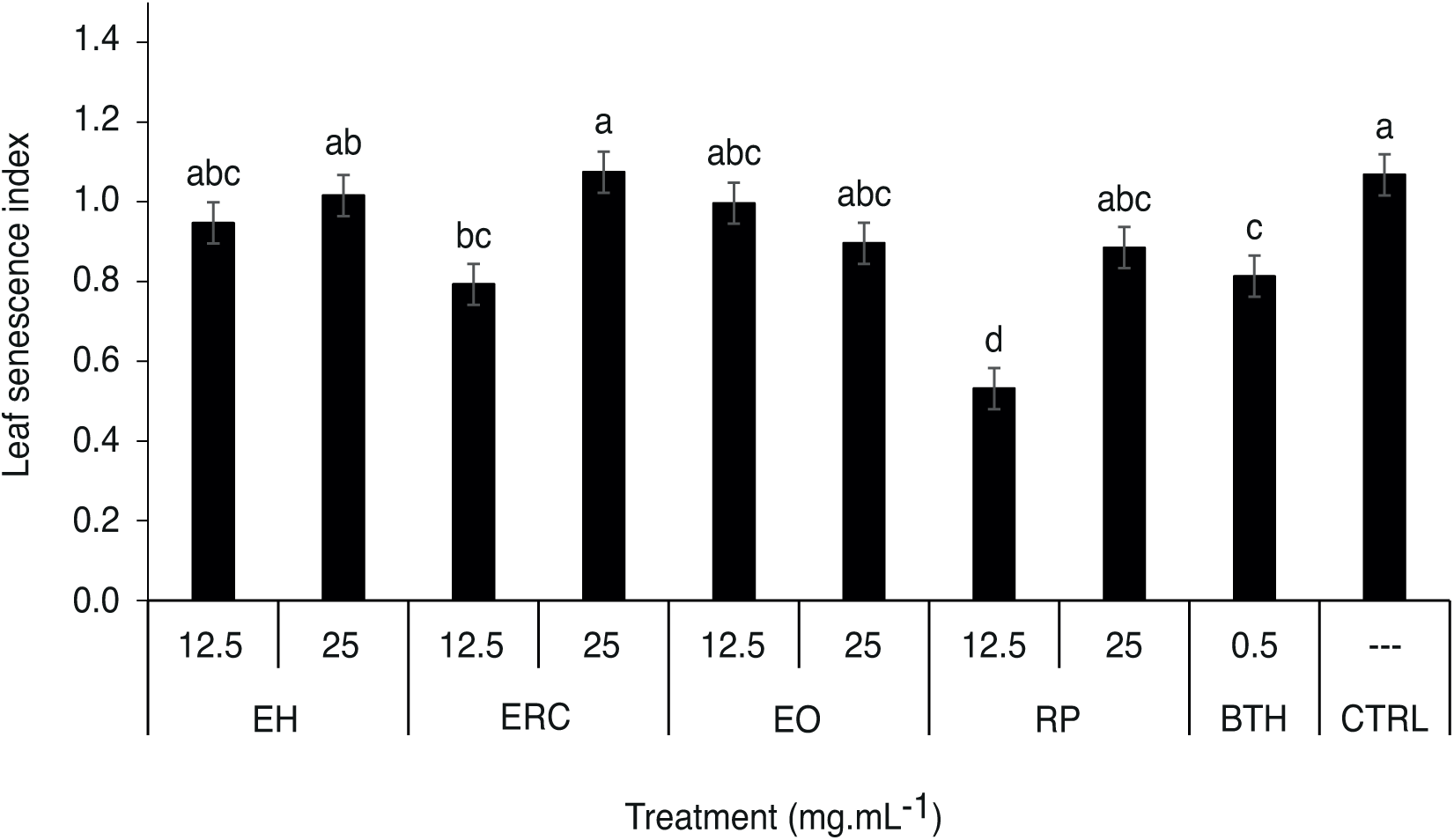
Leaf senescence index for plants treated with Eastern hemlock (EH), eastern red cedar (ERC), English oak (EO) and red pine (RP) extracts at working concentrations of 12.5 and 25 mg mL^−1^, treated with BTH (ACTIGARD^TM^ 50WG) at 0.5 mg mL^−1^ (positive control), or treated with sterile water (negative control). Treatments were applied by foliar spraying (20 mL), two days before plant inoculation with *Pseudomonas syringae* pv. *tomato* DC3000 suspension. Each value is the mean of three biological replicates (with six plants per replicate) ± SE. Mean values with the same letter are not significantly different (post-ANOVA LSD, with an alpha threshold value of 5%).

**Fig. 3.**
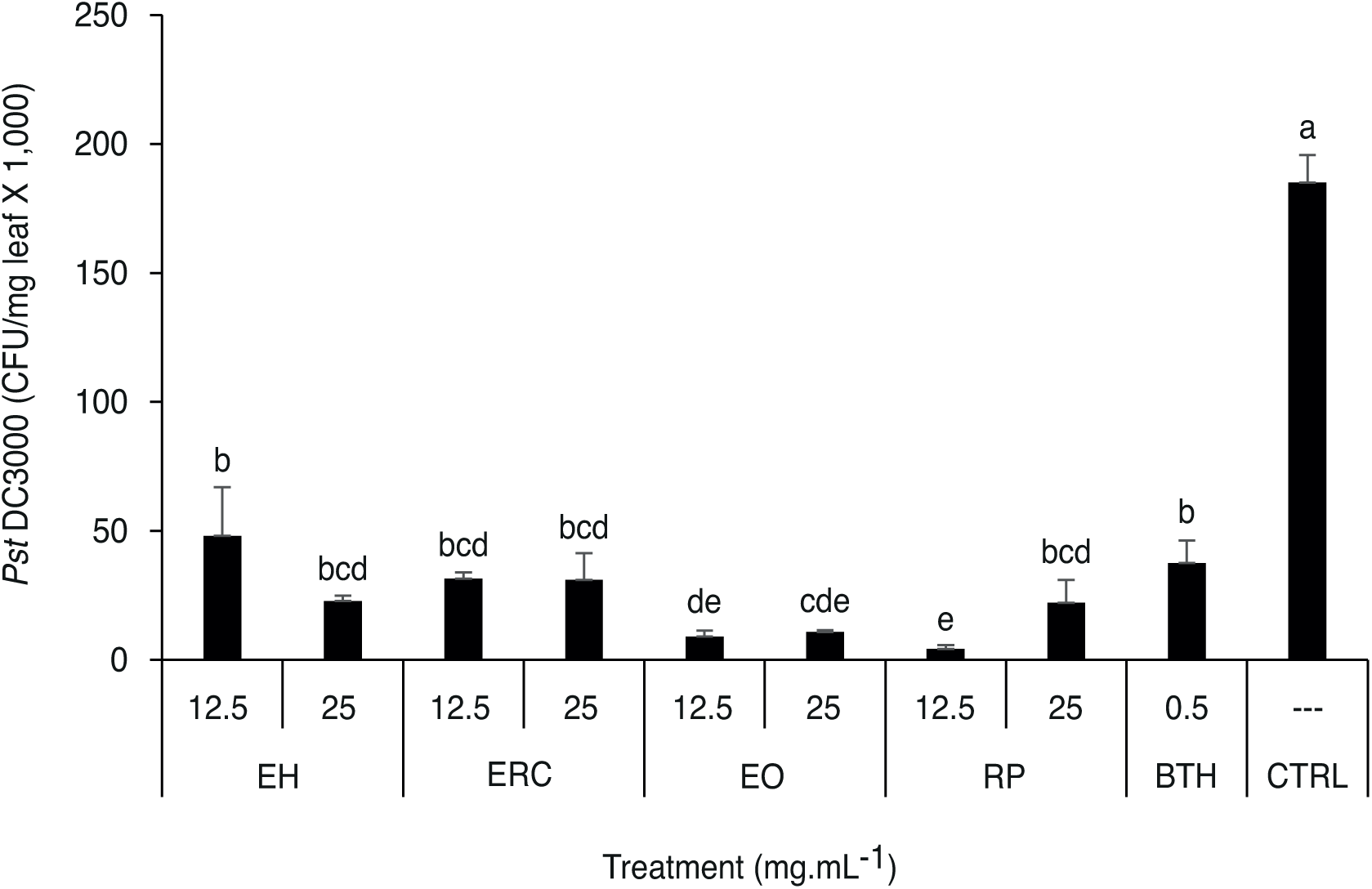
*Pseudomonas syringae* DC3000 populations in leaf tissue of *Arabidopsis thaliana* treated with the eastern hemlock extract (EH), the eastern red cedar extract (ERC), the English oak extract (EO) or the red pine extract (RP) at working concentrations of 12.5 and 25 mg mL^−1^, with BTH (ACTIGARD^TM^ 50WG) at 0.5 mg mL^−1^ (positive control), or with sterile water (negative control). The extracts were applied by foliar spraying (20 mL), two days before plant inoculation with *P. syringae* DC3000 suspension. Each bar is the mean of three biological replicates ± SE. Mean values with the same letter are not significantly different (one-way ANOVA with post hoc LSD with an alpha threshold of 5%).

Leaf senescence indices were first determined to rule out the possibility of phytotoxic effects for the tree extracts and to detect possible senescence-delaying effects for these extracts following bacterial infection, associated with their chemical content and/or respective impacts on ROS production and the induction of ROS-scavenging enzymes in treated leaves (**Fig. 2**). Plants were sprayed with the extracts, two days prior to inoculation with the bacterial pathogen. In brief, senescence indices for the extract-treated plants were similar to, or lower than, the senescence index observed for water-treated plants after 9 days, one-week after plant infection with the bacterial pathogen. Compared to the other plants, those plants treated with the red pine extract showed a leaf senescence index of ∼0.55 at the lowest dose applied, about two times smaller than the leaf senescence index of control plants. A possible explanation for this could involve the SAR inducer WRKY70 in extract-treated plants, given the well-known antagonistic effect of this regulatory protein on leaf senescence [38,68]. An alternative, more likely explanation considering roughly similar induction rates for *WRKY70* by different leaf extracts, would involve the presence of specific chemicals (e.g., terpenoids) in the red pine extract.

Plants treated with the tree extracts were then challenged with *Pst* DC3000 two days post-treatment to assess their overall protective effects against the pathogen and to highlight possible differential effects among the extracts given their variable toxicity against the pathogen and distinct inducing effects on defense- and stress-related genes in leaves (**Fig. 3**). Bacterial populations were quantified one week after leaf inoculation to evaluate the antibacterial potential of the extracts. Bacterial counts of ∼5,000 to 50,000 colony-forming units (CFU) were observed for the extract-treated plants, similar to bacterial counts for BTH-treated plants but far lesser than the bacterial count of ∼200,000 CFU determined for the control plants. The English oak extract, as the most toxic extract against *Pst* DC3000 (**Table 1**) but with a moderate efficiency to induce SAR compared to the eastern hemlock and red cedar extracts (**Table 3**), allowed for a strong, 95% inhibition of bacterial growth compared to the control treatment. The eastern hemlock extract, a potent inducer of SA-inducible marker genes, also allowed for an elevated growth inhibition rate similar to the inhibition rate observed with the non-toxic SA mimic BTH, despite a less pronounced *in vitro* antibacterial effect compared to the other three extracts. Overall, these findings confirmed the practical potential of tree by-product extracts as effective SAR-inducing biopesticides for *P. syringae* control. They also suggested a dual antibacterial effect for the tree extracts, explained in different proportions from one extract to another by direct toxic effects against the pathogen and an effective induction of the host plant natural defenses.

## Conclusion

Several studies have assessed the protective potential of plant extracts against pests and pathogens, as a sustainable, eco-friendly alternative to conventional pesticides. In this study, we showed the potential of by-products from four forest tree species of economic importance in protecting plants from bacterial infection via direct toxic effects on the target pathogen and indirect inducing effects on the host plant’s natural defense system (**Fig. 4**). Complementary studies will be welcome in forthcoming years to identify chemical(s) in the tree extracts that account(s) for the toxic effects against *P. syringae* and other pathogen targets. Studies will also be welcome to elucidate the host plant’s response to the tree extracts. Our data clearly pointed to the establishment of a SAR response in leaf tissue involving the well characterized SAR master regulators WRKY70 and NPR1 and the downstream production of SA-inducible defense compounds, including antimicrobial PR proteins (**Fig. 4A**). Several questions remain regarding the identity of (the) basic chemical trigger(s) of plant defense responses in the extracts, the possible importance of an oxidative stress mitigating response in leaves after extract treatment, the relative roles of WRKY70 and NPR1 in the whole gene inducing process, and the establishment – or not– of an anti-senescence effect upon extract treatment (**Fig. 4B**). Work is underway to address these questions, and to confirm the potential of the four tree extracts for the control of economically relevant microbial diseases.

**Fig. 4.**
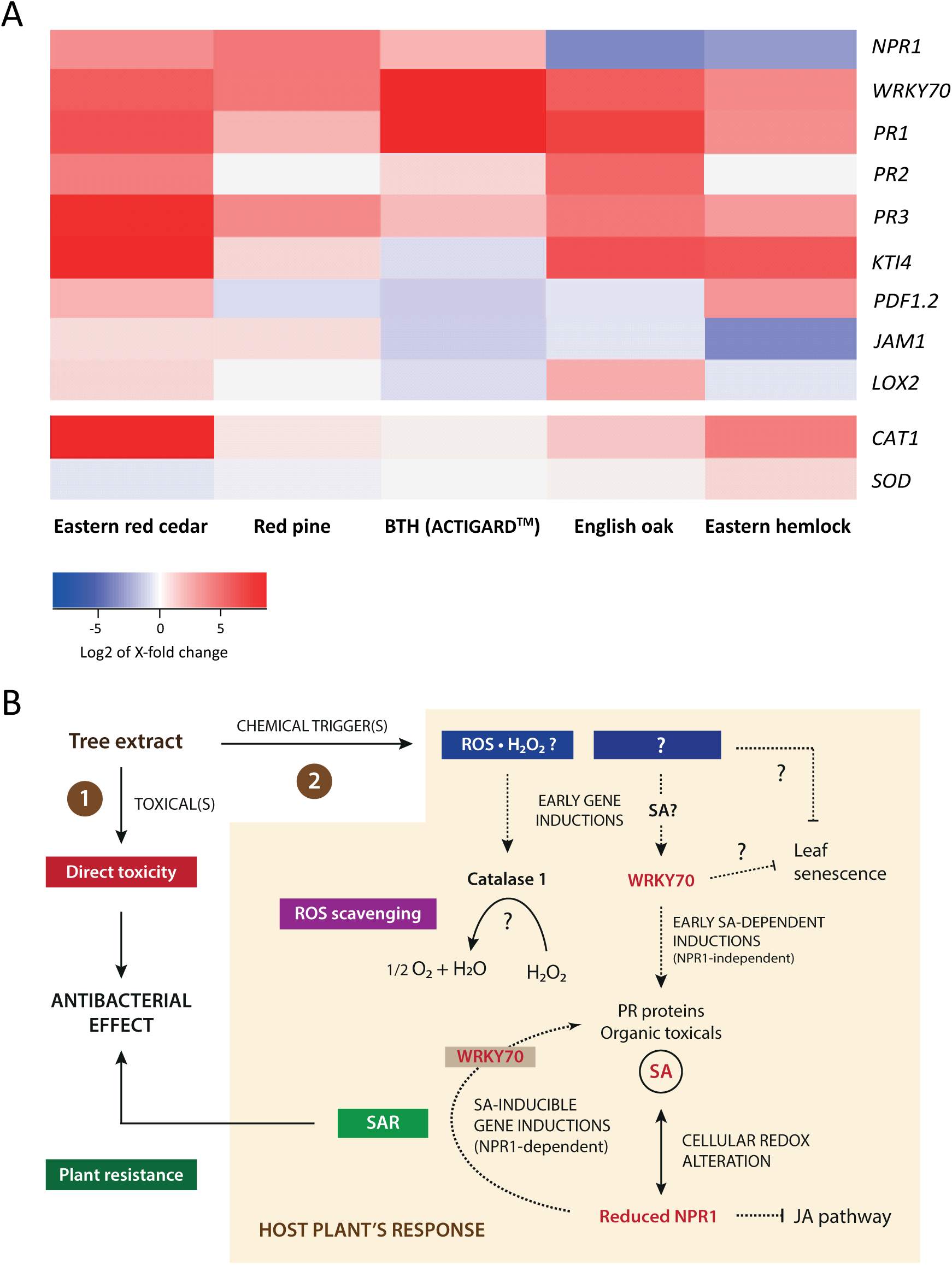
Arabidopsis response to the forest tree extracts – A working frame for future studies. **A,** Heat map for the up- and downregulating effects of extracts on defense and stress marker genes. See Table 2 for genes nomenclature and expected inducibility. Heat maps were produced with log2 values of actual expression ratios reported in Tables 3 and 4. Color intensities correlate with the intensity of the up- and downregulating effects observed. Red boxes highlight gene upregulations, and blue boxes downregulations, compared to control plants. **B,** Working model for the dual impact of the tree extracts on *P. syringae* and Arabidopsis. Extracts exert a dual effect on the bacterial pathogen, involving (**1**) direct toxicity effects dependent on their specific chemical composition (TOXICAL(S)); and (**2**) a defense SAR response conferring the host plant with increased resistance to the pathogen. Several questions remain at this stage regarding the chemical determinants of microbial toxicity in the extracts. Questions also remain about the plant’s response to extract treatments, notably related to the CHEMICAL TRIGGERS of catalase (Cat1) expression for ROS (H_2_O_2_) scavenging induction and EARLY INDUCTION of WRKY70; about the SA-mediated reduction of the cell cytosol milieu required for NPR1 activation leading to SAR [76] and JA pathway inhibition, and about the cause and actual significance of apparent senescence delays observed post-infection following treatment with some extracts. Dotted lines on this panel highlight transcriptional regulations; plain lines highlight chemical reactions or biological effects or responses.

## Methods

### Plant materials and growth chamber conditions

Seeds of transgenic Arabidopsis line PR1::GUS were kindly provided by Prof. Corné M.J. Pieterse (Utrecht University, The Netherlands) [29]. The seeds were surface sterilized and placed on Murashige & Skoog (MS) medium containing 2.22 g L^−1^ Murashige & Skoog Basal salts (GeneLinx International, Inc., DBA bioWorld, Dublin OH, USA), 0.5 g L^−1^ MES hydrate (Acros Organics, Geel, Belgium), 4 g L^−1^ phytagel (Alfa Aesar, Ottawa ON, Canada), and 10 g L^−1^ sucrose (Anachemia Inc., Montréal QC, Canada) (adjusted to pH 5.7). The Petri dishes were incubated for two days at 4 °C, and then placed vertically in a growth chamber for 14 days, to allow for the roots to grow on the surface. The plants were kept under a light intensity of 80 μmol photons m^−2^ s^−1^ during the day, a light photoperiod of 16 h and a temperature regime of 23 °C during the day and 18 °C during the night. For *in vivo* assays, the seedlinga were transplanted in 6-cell seedling starter trays containing Veranda Mix Potting Soil (Scotts Fafard, Saint-Bonaventure QC, Canada), and the plants grown on soil in the controlled growth chamber for 40 days, under a 16 h light photoperiod at 21 °C, a light intensity of 210 μmol photons m^−2^ s^−1^ and a relative humidity of 70%. Plants were irrigated as needed and fertilized twice a week with a 50 ppm NPK (20:20:20) solution.

### Tree by-product extracts

Leaves of English oak and twigs of eastern hemlock, eastern red cedar and red pine were collected at the Jardin universitaire Roger-Van den Hende (Université Laval, Québec, QC, Canada). The crude extracts were prepared as described by Delisle-Houde et al. [18]. In brief, the tree samples were ground mechanically and macerated under agitation (100 rpm) for 24 h in 95% (v/v) ethanol at 22.5 °C. Each extract suspension was then evaporated at a temperature below 60 °C using a R-200 rotary evaporator (Büchi Labortechnik AG, Flawil, Switzerland). The resulting powder was freeze-dried and stored in the dark at room temperature in Mason jars.

### *Pst* DC3000

Rifampicin-resistant *Pst* DC3000 was used for the experiments [69]. The bacteria were kept in 15% (v/v) glycerol (VWR International, West Chester PA, USA) diluted in water until use, and cultivated at 28 °C on King’s B (KB) nutrient [20 g L^−1^ of Bacto™ Proteose Peptone No. 3 (Becton, Dickinson, and Company, Sparks MD, USA) supplemented with 1.5 g L^−1^ dibasic sodium phosphate (EM Science, Gibbstown NJ, USA), 10 g L^−1^ glycerol (VWR International), and 1.5 g L^−1^ magnesium sulfate (Fisher Scientific, Geel, Belgium) solid medium [15 g L^−1^ of CRITERION™ Agar (Hardy Diagnostics, Santa Maria CA, USA)]. The bacteria were then inoculated in flasks containing 20 mL of KB nutrient liquid medium, allowed to grow under agitation (160 rpm) for 24 h at 28 °C, recovered by centrifugation for 5 min at 3600 × g, and finally suspended in 10 mM MgSO_4_ (Fisher Scientific) aqueous solution at 1 × 10^8^ CFU mL^−1^. The suspension was supplemented with 0.01% (w/v) SYLGARD™ OFX-0309 (Dow Chemical Company, Midland MI, USA) for the growth chamber assays. Bacterial concentrations were monitored by measurement of optical density at 600 nm (OD_600_), using 0.5 McFarland standards and an Epoch 2 Microplate Spectrophotometer (BioTek Instruments, Inc., Winooski, VT, USA).

### Minimum inhibitory concentrations

Minimum inhibitory concentrations (MICs) were determined under sterile conditions against *Pst* DC3000 using flat-bottom 96-well microplates (Sarstedt AG & Co., Nümbrecht, Germany). Bacterial cells (at 5 × 10^5^ CFU) were suspended in 100 μL of KB liquid nutrient medium containing different concentrations of each plant extract (0, 0.39, 0.78, 1.56, 3.13, 6.25, 12.5, 25, and 50 mg mL^−1^). The microplates were incubated for 24 h at 28 °C, and 10 μL of sterile 2,3,5-triphenyl-2H-tetrazolium chloride (1 mg mL^−1^; Ward’s Science, Rochester NY, USA) was then added to each well [70]. For each plant extract, the lowest concentration at which no metabolic activity was observed (absence of red coloration in the well) corresponded to the MIC. Each concentration was tested in three replicates and the experiment was conducted twice.

### Minimum bactericidal concentrations

Minimum bactericidal concentrations (MBCs) were determined under sterile conditions using flat-bottom 96-well microplates as described above for the MICs, except that the content of each well was spread on KB solid medium in Petri plates after 24 h incubation, instead of adding 2,3,5-triphenyl-2H-tetrazolium chloride [70]. The Petri plates were incubated for 48 h at 28 °C. For each plant extract, the lowest concentration at which no growth was observed on KB solid medium corresponded to the MBC. Each concentration was tested in three replicates and the experiment was conducted twice.

### Activation of host plant defenses

Ethanolic plant extracts cold sterilized with 0.22 μm filter papers were added (5 mL) to Petri dishes containing 14-day old seedlings of Arabidopsis transgenic line PR1::GUS at two concentrations (12.5 and 25 mg mL^-1^). Milli-Q water and BTH (ACTIGARD^TM^ 50WG; Syngenta Canada Inc., Guelph, ON, Canada; 0.5 mg mL^−1^) in sterile water were used as negative and positive controls, respectively. The Petri dishes were incubated for 48 h at 22.5 °C, and the seedlings then submitted to a GUS activity assay as described below. Quantitative RT-PCR assays were also performed to assess the activation of defense-related genes in the treated plants, using DNA primers for different defense gene markers (**Table 5**).

**Table 5.**
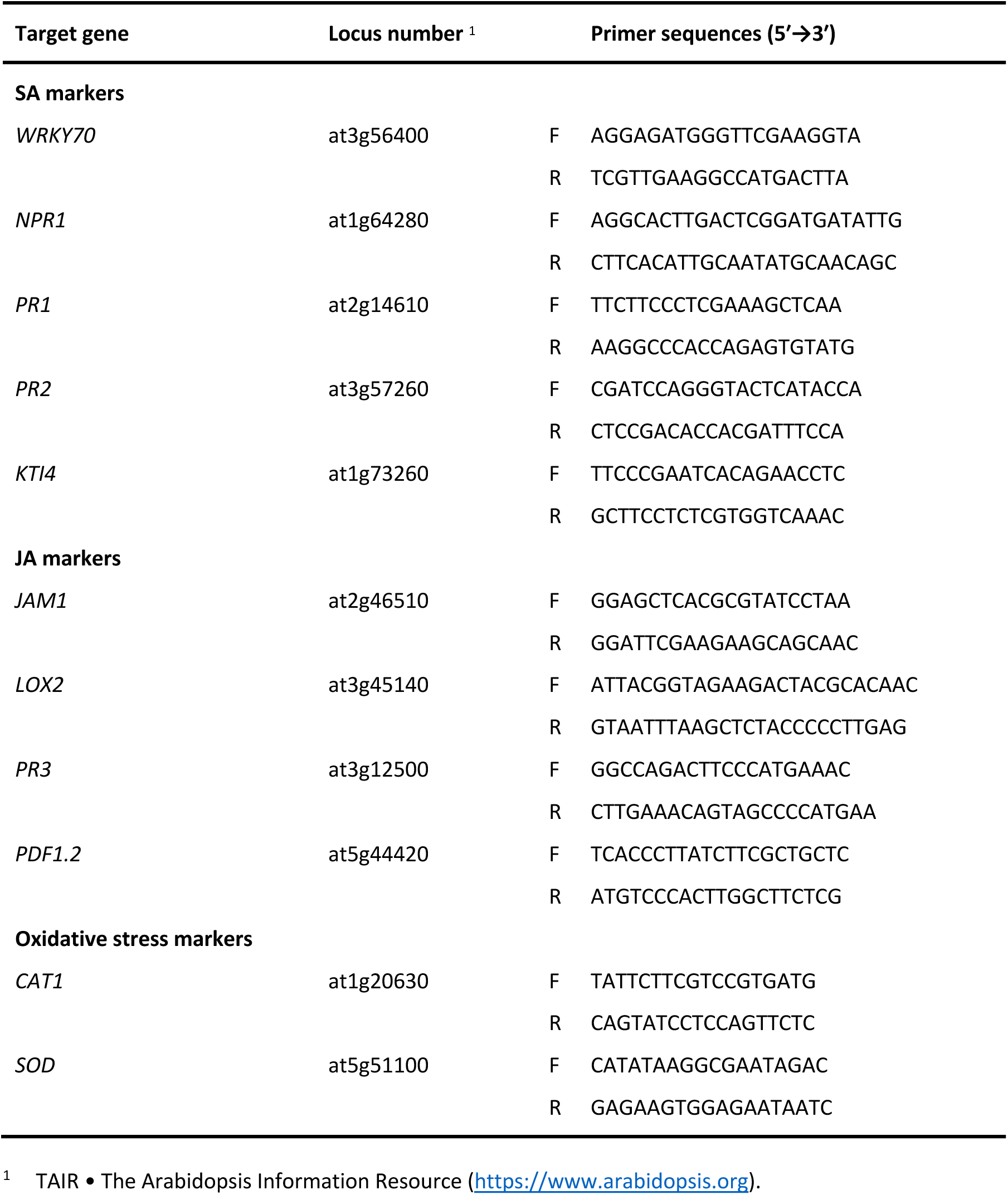
Primers used for the RT-qPCR analyses.

### GUS activity assay for *PR1* promoter induction

Seedlings or leaves from blooming stage plants of Arabidopsis PR1::GUS line were transferred in a microtube containing 2 mL of GUS staining solution [1 mM X-Gluc (Biosynth International, Inc., San Diego CA, USA), 100 µM phosphate buffer (Fisher Scientific, Hampton NH, USA), 10 mM EDTA (Fisher Scientific), 0.1% Triton (v/v) (Fisher Scientific), 1 mM potassium ferrocyanide (II) (Sigma-Aldrich Inc., St. Louis MO, USA) and 1 mM potassium ferricyanide (III) (Sigma-Aldrich Inc.)] for 16 h at 37 °C. The samples were washed one time with 70% (v/v) ethanol to remove the GUS staining solution, incubated for 1 h in 70% (v/v) ethanol at 60 °C to extract chlorophyll from plant tissues, and washed three times in ethanol 70% (v/v) to remove the pigment. The leaves and seedlings were subjected to microscopy observation using a SZ2-ILST Olympus stereomicroscope (Olympus Co. Tokyo, Japan). Images were assembled using the photo merge tool of Adobe Photoshop CS6 (Adobe, Mountain View CA, USA). GUS staining was quantified using the image analysis software ImageJ [71].

### RNA isolation and RT-qPCR

RNA extraction and RT-qPCR were performed as described in Barrada et al. [72]. Seedlings (n = 10) of Arabidopsis *PR1::GUS* line from the *in vitro* assay were taken 48 h after the treatments and ground to a fine powder in liquid nitrogen. Total RNA was extracted using the Rapid Plant RNA Isolation Kit (Bio Basic, Markham ON, Canada). RNA quality was confirmed by gel electrophoresis, and genomic DNA removed by treatment with DNase I (Thermo Fisher Scientific Inc., Waltham MA, USA). cDNA was synthesized from 500 ng RNA using a Primescript RT Reagent Kit (Takara Bio, Shiga, Japan) with random hexamer and oligo(dT) primers, and stored at -20 °C until use. Quantitative RT-PCR was performed in 1 μL of one-fifth diluted cDNA in 35-μL reactions containing the SYBR™ Green I Nucleic Acid Gel Stain (1:20,000 final dilution; Thermo Fisher Scientific), Taq DNA polymerase (1.75 units) with standard Taq buffer (New England Biolabs, Pickering ON, Canada), 0.2 mM dNTPs and 0.2 μM primers (**Table 1**). The PCR reactions were performed in a LightCycler96® instrument (Roche, Basel, Switzerland) real-time system under the following conditions: an initial denaturation cycle of 30 s at 95 °C, followed by 45 cycles of denaturation (10 s at 95 °C), annealing (30 s at 60 °C), and polymerization (30 s 72 °C). Relative quantification of gene expression adjusted for efficiency was performed using PCR Miner [73]. *Actin2* (*ACT2*), *glyceraldehyde-3-phosphate dehydrogenase* (*GAPDH*), *18S-rRNA*, and *WRKY1* were used as reference genes. The stability of reference genes relative expression was validated according to Vandesompele et al. [74], with advised M and Cv limits of < 0.5 and < 0.25, respectively.

### Effect of the extracts on leaf senescence and *Pst* DC3000 populations

During the growth chamber assay, plant extract suspensions, BTH solution (20 mL per plant) or milli Q water were added by foliar spraying to Arabidopsis lines PR1::GUS cultivated in a growth chamber under the conditions described above. Plant extracts were used at the same concentrations as for the *in vitro* assay on plant at the blooming stage. The formulations were sprayed on plants two days before inoculation with a suspension of rifampicin-resistant *Pst* DC3000 mutant at 1 × 10^8^ CFU mL^−1^.

Leaf senescence was evaluated for each plant using a leaf senescence index determined according to **Equation 1**. The experiment was conducted as a completely randomized design, with two replicates each including six plants (n = 12).

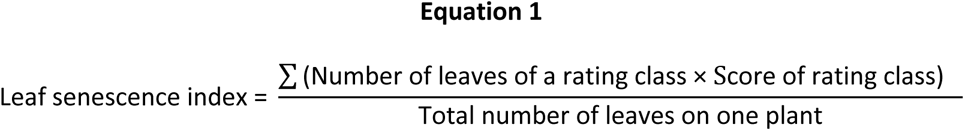

where the rating class corresponded to the extent of affected leaf area, based on the following categories : 1 = less than 25%, 2 = 26% to 50%, 3 = 51% to 75%, and 4 = 76% to 100%.

One week after inoculation, viable bacterial (*Pst* DC3000) populations were determined in plant tissues according to a protocol adapted from Jacob et al. [75]. For each treatment, three leaves of similar size and age (representing 200-300 mg) were taken randomly on three different plants in each 6-cell seedling starter tray and ground two times (30 s, in 1 mL of 10 mM MgSO_4_ in water), using an Omni Bead Ruptor Homogenizer (Omni International Inc., Kennesaw GA, USA). Ten microliters of each suspension were serially diluted and 10 μL of each dilution were spread on KB solid medium in Petri plates supplemented with rifampicin (Sigma-Aldrich Inc.) at 50 mg L^−1^. After incubation for 30 h at 28 °C, the number of CFU was counted. Populations of *Pst* DC3000 were expressed as CFU per milligram of fresh leaf.

### Statistical analyses

Analyses of variance (ANOVAs) and Chi-square tests were performed on the data using the R software (R-4.1.1, R Core Team, 2021, Vienna, Austria). Raw data were transformed using Tukey’s ladder of powers, and the treatments then compared using a *post hoc* LSD test. A non-parametric Kruskal–Wallis test (p ≤ 0.05) was performed in parallel in R. The regular parametric approach was deemed valid if it gave results nearly identical to the non-parametric approach [16].

## Acknowledgements

Authors extend their heartfelt gratitude to Prof. Corné M.J. Pieterse (Utrecht University, Utrecht, The Netherlands) for providing seeds of transgenic Arabidopsis PR1::GUS line, and to Dr. Edel Pérez López (Université Laval, Québec QC, Canada) for providing bacterial strain *Pst* DC3000.

## Authors’ contributions

Conceptualization: V.S., A.B., M.D.H., R.J.T., D.M.; data acquisition: V.S., A.B. M.D.H.; data analysis: V.S., A.B., M.D.H.; data interpretation: V.S., A.B., M.D.H., R.J.T., M.D., D.M.; draft manuscript preparation: V.S., A.B., M.D.H., R.J.T., D.M. All authors reviewed and approved content of the manuscript before submission.

## Data availability

The data presented in this study are available on request from the corresponding authors.

## Declarations

### Ethics approval and consent to participate

All experiments were conducted in accordance with relevant institutional, national, and international guidelines and legislation.

### Consent for publication

All authors consent to the publication of the manuscript.

